# Autophagy, the Ubiquitin proteasome system, and the MAPK pathway control the temperature dependence of synaptic growth

**DOI:** 10.1101/2024.10.03.616550

**Authors:** Kevin De Leon, Bruno Marie

## Abstract

There is clear evidence that Earth’s temperature is rising at an unprecedented rate. While consequences on ecosystems are being extensively studied, little is known about the consequences of temperature on the nervous system of ectothermic animals. Here we used the *Drosophila* model NMJ to assess the effect of temperature on the synaptic growth of a phasic, and a tonic, motoneuron. We find that the tonic neuron’s synaptic size is not affected by temperature, however we do observe a temperature-dependent synaptic growth in the phasic neuron, which might be related to the increased motility observed previously at higher temperatures. We find that the level of autophagy activity changes with temperature and that autophagy genes are responsible for the temperature dependence of synaptic growth. We present evidence that this regulation could occur through the major synaptic growth regulator and ubiquitin ligase Highwire, and a pathway involving the Mitogen-Activated Protein Kinases. We present a new function for the MAPKKK, Wallenda and the MAPK P38b in directing the additional synaptic growth that takes place between 25°C and 29°C. This illustrates that temperature has different effects on a diverse population of neurons and that distinct genetic pathways are involved in regulating temperature driven changes.

## Introduction

Temperature is a major environmental factor able to affect nervous systems (Andersen & Moser, 1995; Hodgkin et al., 1952; Kim et al., 2022; Micheva & Smith, 2005). These effects are present in endothermic animals, where slight changes in temperature within the mammalian nervous system can affect synaptic/neuronal properties (Kiyatkin, 2010; H. Wang et al., 2014). Temperature changes can affect ectothermic animals, such as fruit flies, more profoundly because they can be long term and affect developmental processes. In *Drosophila* temperature can influence synaptic growth (Kiral et al., 2021; Mellert et al., 2016; Peng et al., 2007; Sigrist et al., 2003) and activity-dependent synaptic plasticity (Chopra & Singh, 1994; Kiral et al., 2021; Maldonado-Díaz et al., 2021; Sigrist et al., 2003; Yeates et al., 2017). There has been a broad interest in understanding the consequences of extreme temperature in the nervous system (cold shock and heat shock) (Al-Fageeh & Smales, 2006; Micheva & Smith, 2005; Parsell & Lindquist, 1993; Phadtare et al., 1999; Velazquez & Lindquist, 1984) but, in a warming planet (IPCC, 2022; IPCC, 2023), there is also an increasing interest in understanding how smaller changes in temperature can affect the function of the nervous systems (Kushmerick et al., 2006; Owen et al., 2019; H. Wang et al., 2014). To date, one of the most noticeable characterizations of the effect of temperature on the nervous system remains the example of the Drosophila larval neuromuscular junction (NMJ; (Sigrist et al., 2003). In this study, larvae raised at higher temperatures (29°C) showed an increase in synaptic growth compared to larvae reared at 18°C or 25°C.

Because differences in temperature can affect several aspects of the biology of ectothermic animals, it is tempting to relate temperature-dependent changes to major physiological processes like metabolism and activity/locomotion, which in turn could have multiple phenotypic expressions (Kiral et al., 2021; Li & Gong, 2015; Sigrist et al., 2003). This should not exclude the possibility that the changes in synaptic size, consequent to changes in temperature, involve discrete genetics pathways regulating the temperature-dependence of synaptic growth. Indeed, the NMJ has been a model of choice to understand the molecular mechanisms governing synaptic growth (Budnik & Ruiz-Canada, 2006; Collins & DiAntonio, 2007; Keshishian et al., 1996; Van Vactor & Sigrist, 2017) and the idea that molecular determinants could differentially respond to temperatures has yet to be tested.

In addition, the nervous system is composed of a great diversity of neurons and the question remains whether some neurons might be more susceptible to temperature than others. Is there a plurality of molecular mechanisms in the neuronal response(s) after temperature changes? The Drosophila model NMJ can allow us to test this question. Indeed, because it is stereotypical and easily accessible, the synapse innervating muscles 6 and 7 has been extensively characterized (Johansen et al., 1989; Keshishian et al., 1996). But it is important to note that this glutamatergic synapse consists of two different terminal projections (Is and Ib) originating from a phasic and a tonic motoneuron, respectively (Aponte-Santiago et al., 2020; Atwood et al., 1993; Jia et al., 1993; Lnenicka & Keshishian, 2000; Schuster et al., 1996). Lately, evidence has emerged characterizing the numerous differences between these two motoneurons regarding their anatomy, physiology, plasticity and their differentially expressed genes (Aponte-Santiago et al., 2020; He et al., 2023; Hoang & Chiba, 2001; Jetti et al., 2023; Lu et al., 2016; Newman et al., 2017; Nguyen & Stewart, 2016; Y. Wang et al., 2021).

Here we not only reiterate the experiments showing the temperature dependence of synaptic growth using temperatures spanning from 15°C to 29°C, but we also showed that temperature only affects the Is synapses, with the Ib synapses being temperature-independent. We then asked whether genes known to control synapse size were involved in temperature-dependent growth. We found that the autophagy gene *atg1* controls temperature-dependent synaptic growth and that the endogenous autophagy activity varies with the temperature, suggesting that temperature could affect autophagy and therefore synaptic size. In addition, we characterized a known regulator of synaptic growth, the E3 ubiquitin ligase Highwire and show that it could affect both the temperature-dependent growth of the Is boutons and the temperature-independent growth of the Ib motoneurons. Finally, we found a novel role for MAP Kinase signaling, in which Wallenda (MAPKKK) and P38b (MAPK) are necessary to promote synaptic growth in Is motoneurons only at higher rearing temperatures.

## Material and Methods

### Genetics

Animals of either sex were used in this study. All animal lines were obtained from Bloomington Stock Center. *w^1118^* (BDSC stock #145) was used as control strain. To test autophagy loss-of-function mutants, we used an allelic combination of *atg1^Ey09216^*(BDSC stock #16816) and *atg1^3^* (BDSC stock #60732). For the loss of function of the E3 ubiquitin ligase Highwire, we used *hiw^N^*(BDSC stock #51637). To test the loss of function of MAPKKK and MAPK Wallenda and P38b we used the alleles *wnd^3^* (BDSC stock #51999), *wnd^1^*(BDSC stock #51641) and *p38b^KG01337^*(BDSC stock #14364). In addition, we used *UAS-hep^Rnai^* (BDSC stock #35210), *UAS-JNK^DN^* (BDSC stock #9311), and *UAS-JNK^Rnai^* (BDSC stock #57035), *mkk4^e01485^* (BDSC stock #35143) and *p38a^1^* (BDSC stock #8822).

### Rearing methods

All animals were reared in standard *Drosophila* cornmeal media, Jazz-mix (Fisher Scientific, #AS153). The animals were reared in temperature-controlled incubators at temperatures of 15°C, 25°C, and 29°C. Experiments for synaptic growth at the neuromuscular junction were performed on animals at the wandering third instar larval stage.

The rearing of second instar larvae animals at 15°C and 29°C (Ref(2)P and Lysotracker quantification experiments) were performed on apple juice agar plates supplemented with yeast paste. The CNS of early second instar larvae was collected for the experiments. We distinguished the larval stage by the size and presence of visible mouth hooks.

### Immunohistochemistry

For experiments examining synaptic growth, third-instar larvae were dissected in HL3 saline (70mM NaCl, 10 mM NaHCO3, 115 mM sucrose, 5 mM treha-lose, 5 mM HEPES, 10 mM MgCl2, 5 mM KCl and 0.1 mM CaCl2) and fixed in Bouin’s fixative (Sigma) for 1 min. For experiments involving the anti-Ref(2)P antibody the muscle and CNS of the larval preparations were fixed in

4% paraformaldehyde for 15 min. After fixation preparations were washed in PBT 0.1% for 1 hour. Primary antibodies were used overnight at 4°C. We used a newly made affinity-purified rabbit anti-Dlg antibody [1:200] (PrimmBiotech, Inc.) raised against a recombinant protein containing the Dlg sequence stretching from amino acid 764 to amino acid 919. This polyclonal rabbit antibody provided immunoreactivity identical to the monoclonal mouse anti-Dlg (Developmental Studies Hybridoma Bank, DSHB). We also used the mouse anti-Syn, [1:20, (DSHB)] and the rabbit anti-Ref(2)P [1:200, Abcam (ab#178440)]. Secondary antibodies and the anti-HRP antibody were applied for 1 hour at room temperature as previously described (Maldonado et al., 2013; Marie et al., 2004). These antibodies were AffiniPure anti-HRP (Jackson ImmunoResearch) conjugated to Cy3 (1:300) and the secondary antibodies Alexa Fluor 488-conjugated AffiniPure goat anti-mouse (1:300) and Alexa Fluor 545-conjugated AffiniPure goat anti-rabbit (1:300). The preparations were mounted in Vectashield (Vector Laboratories) for microscopic analysis.

### Quantification of synapse size

To quantify synaptic growth, we scored m6/7 synapses in segment A3 of third-instar larvae. Synaptic boutons were revealed by anti-Synapsin and anti-Dlg immunolabeling. The Is and Ib motoneurons were distinguished by the intensity of DLG staining and were counted and averaged using anti-Synapsin staining. We present in the figures each specific genotype and condition, the average, SEM and scatter plot derived from a minimum of 8 synapses from at least 4 animals. We used a Nikon Eclipse 80i microscope at a magnification of 40X to quantify synaptic boutons. The images were acquired on a Nikon A1R resonant scanning confocal microscope using a 40X objective. We obtained a series of optical sections at 0.2 µm intervals encompassing the whole synapse delineated by HRP staining. Maximum intensity projections were obtained, and composite images were converted into Tiff format using the open-source ImageJ Java-based image processing and analysis program (National Institutes of Health; http://imagej.nih.gov/ij/). These images were assembled, and the contrast of the figures was adjusted using Photoshop CC2018 (Adobe Systems). Charts were created using Prism 6 (GraphPad).

### Lysotracker fluorescence

The experimental design was adapted from Shen and Ganetzky (2009). In summary, brains from second instar larvae of control animals reared at 15°C or 29°C were dissected in HL3 saline, incubated for 5 min in 0.1 µM Lysotracker red DND-99 (Invitrogen) in HL3 saline, rinsed twice in PBS (Sigma#P4417, 10 mM phosphate buffer, 2.7 mM potassium chloride and 137 mM sodium chloride), transferred to glass slides coated with poly D-lysine-coated slides (Sigma A-003-E), and imaged immediately using a Nikon A1R resonant scanning confocal microscope with a 40X objective. All samples were processed and imaged under identical conditions.

### Quantification of fluorescence intensity (Ref(2)P and Lysotracker)

For quantification of fluorescence intensities, the preparation was processed and imaged identically using a Nikon A1R resonant scanning confocal microscope with a 40x objective. The whole CNS of second-instar larvae was scanned with a Z-stack of 0.2 µm intervals, and a 2D maximum intensity Z projection was made. The entire brain lobe area was selected using ImageJ software (https://imagej.nih.gov/ij/) and the average fluorescence was calculated. The area outside the CNS was selected and quantified to determine the background intensity level. The fluorescence intensity values represent the difference between the brain lobe intensity and background intensity (ΔF) over the intensity of the background (F) normalized to wild-type values (Marie et al., 2004, 2010; Maldonado et al., 2013).

### Statistical treatment

We first assessed whether data was normally distributed by performing a Shapiro-Wilk normality test (GraphPad Prism 6). When the sample distribution was normal, we ran a parametric one-way ANOVA when comparing more than two samples. The *post hoc* Tukey’s multiple comparisons test was used for multiple comparisons between data sets. When only two data sets were compared, we performed an unpaired, two-tailed t test. When the Shapiro– Wilk normality test was low (p < 0.01), we ran a nonparametric test. To compare two samples, a Mann-Whitney analysis was performed. The results of these statistical treatments are shown in the graphs of the different figures, and the specific test used is described in the figure legend.

## Results

### The temperature dependence of synaptic growth affects the phasic but not the tonic motoneuron at the *Drosophila* NMJ

We first revisited the data assessing the synaptic growth at different temperatures, extending their range (15°C, 25°C, 29°C). We raised the animals at these different temperatures and asked whether their synaptic sizes were different. We used anti-HRP and anti-Synapsin immunofluorescence to visualize and quantify the number of synaptic boutons at m6/7 segment A3 synapses (see Materials and Methods; Alicea et al., 2017). As previously described, we found that there are clear differences between the mean bouton numbers of wild type NMJs from animals raised at different temperatures (Sigrist et al., 2003). The synaptic size increases with the temperature. Indeed, synapses of animals raised at 15°C have a mean bouton number of 53.4 ± 1.2 (n = 23; Fig.1 A-D, J) while it is 73.9 ± 1.2 (n = 21; p ˂ 0.0001; Fig.1 J) for animals raised at 25°C and 94.7 ± 1.8 for animals raised at 29°C (n= 26, p ˂ 0.0001; Fig.1 G-J).

**Figure 1:**
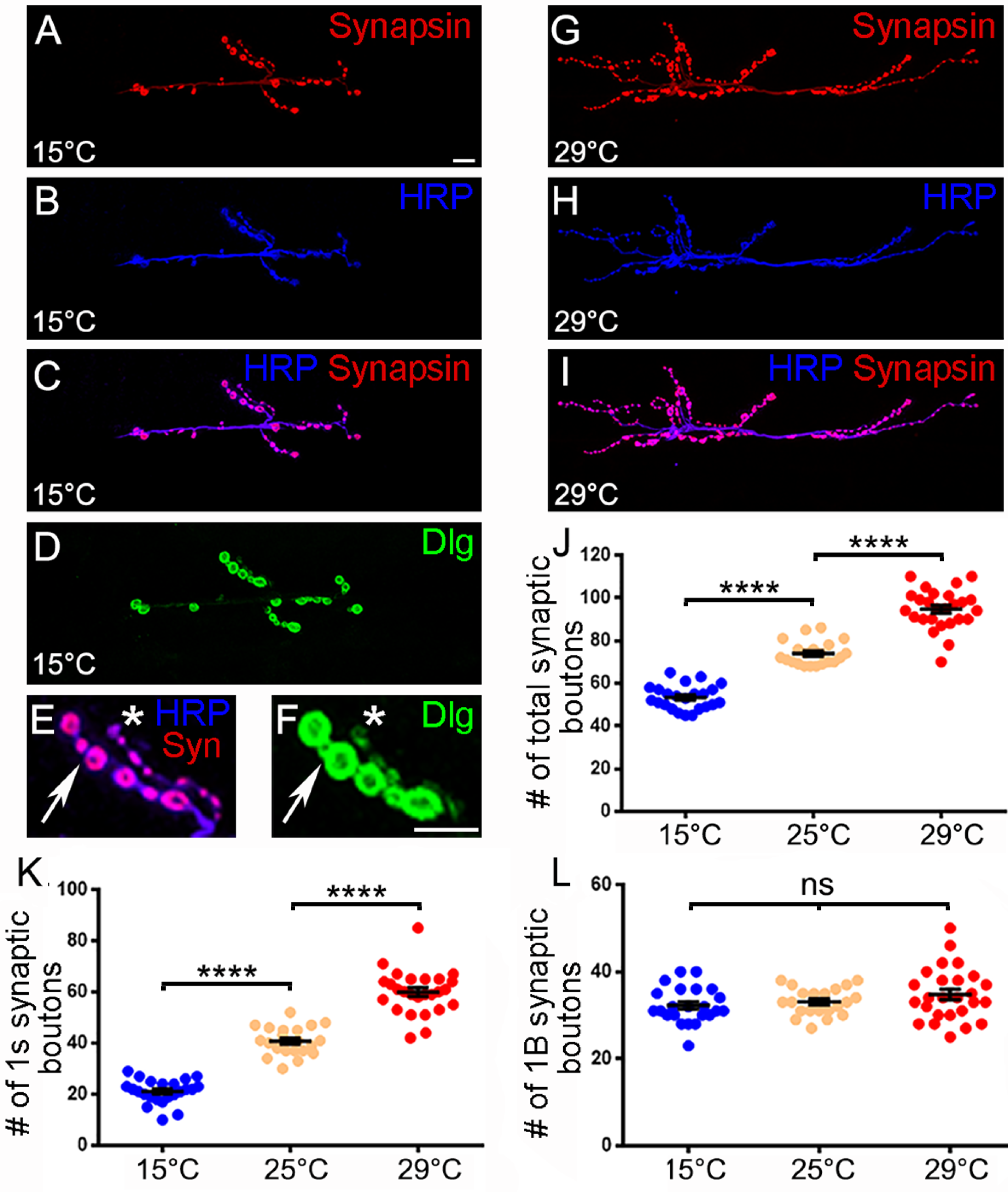
The temperature dependence of synaptic growth is specific to the Is synapses. A-C and G-I: Representative NMJs at m6/7 segment A3 of animals reared at 15°C or 29°C with immunofluorescence for a presynaptic membrane marker (blue; anti-HRP) and presynaptic vesicle marker (red; anti-Synapsin). D: NMJs at m6/7 segment A3 of showing immunoreactivity for the post synaptic marker Discs-Large (green; anti-DLG). E and F: Higher magnification view of part of the NMJ showing anti-HRP, -Syn and -Dlg immunoreactivity and differentiating between the Is (star) and Ib (arrow) synapses. J: Quantification of the total number of synaptic boutons at different temperatures. K: Quantification of the number of the Is synaptic boutons at different temperatures. L: Quantification of the Ib synaptic boutons at different temperatures. **** is p < 0.0001; ns is not significant. ANOVA with post hoc Tukey’s multiple comparisons test. All quantifications show SEM. Scale is 10 μm.

Interestingly, the Drosophila m6/7 model NMJ is composed of two synaptic terminals originating from two different motoneurons. Studies have shown that these motoneurons possess different electrophysiological properties and show differences in genes expression and synaptic morphologies (Aponte-Santiago et al., 2020; He et al., 2023; Jetti et al., 2023; Lu et al., 2016; Newman et al., 2017; Nguyen & Stewart, 2016; Y. Wang et al., 2021). This latter property allows the differentiation of the two terminals. Indeed, one terminal is composed of small synaptic boutons (referred as “Is boutons”, see star in Fig.1 E, F) showing little expression of the postsynaptic Dlg marker [the Cask homologue (Guan et al., 1996)]. A second motoneuron forms on the muscle a terminal composed of larger boutons (referred as “Ib” boutons) and is rich in DLG (see arrow in Fig.1 E, F). We took this opportunity to examine the effect of temperature on these two different motoneurons. We found that temperature seems to exclusively affect the Is synapse (21.1 ± 1 boutons at 15°C, 40.8 ± 1.2 at 25°C and 60 ± 1.7 at 29°C; p < 0.0001; Fig.1 L). To our surprise there is no effect of temperature on the Ib boutons; the mean bouton number is 32.3 ± 0.9 at 15°C; 33.1 ± 0.7 at 25°C and 34.8 ± 1.2 at 29°C (Fig.1 K). While we cannot rule out that the Ib motoneuron could be affected differently by temperature, these data shows that there is a clear effect of temperature on synaptic growth and that this effect is specific to the Is motoneurons. This result exemplifies the diversity in the neuronal response to temperature and suggests that the temperature dependence of synaptic growth is not only the consequence of a ubiquitous physiological process (eg. increase in metabolism) but might be specifically affecting specific neurons and, within these neurons, a particular set of molecular mechanisms regulating synaptic growth. We consequently decided to assess known synaptic growth regulators molecules for their ability to affect temperature-dependent synaptic growth.

### Autophagy controls the temperature dependence of synaptic growth

We decided to focus on the role of autophagy in controlling this process. Indeed, autophagy has been characterized as a positive regulator of synaptic growth (Bhukel et al., 2019; Compans et al., 2021; Kiral et al., 2020; Shen & Ganetzky, 2009; Stavoe et al., 2016). In Drosophila, mutations of the *atg1*, *atg2*, *atg6* and *atg18* genes lead to undergrown synapses (Shen & Ganetzky, 2009). We asked whether *atg1* mutant animals would affect the temperature dependence of synaptic growth. We reasoned that, if autophagy affects synaptic growth but without affecting its temperature dependence, we would expect to observe an effect of temperature on the synapse (smaller at 15°C, higher at 29°C; Fig.2 E, left schematic graph). On the contrary, if the *atg1* mutation does affect the temperature dependence of synaptic growth the mean number of synaptic boutons will be equally undergrown at every temperature (Fig.2 E, right schematic graph). When we carried out the experiment, we observed the latter scenario. The *atg1* mutant animals presented similar undergrown synapses at all temperatures. Indeed, while we observe a temperature-dependent growth at control synapses (52.6 ± 1.5 boutons at 15°C, 73.3 ± 1.6 at 25°C and 97.7 ± 3.1 at 29°C; p < 0.0001; Fig.2 A, B and F), the mean bouton number in *atg1* mutant animals is constant at all temperatures. At 15°C the synapse presents 48.4 ± 2.7 boutons; at 25°C, 49.5 ± 2.6; and at 29°C, 57.4 ± 2.3 (Fig.2 C, D, F). Contrary to what is observed in control animals, raising the *atg1* mutant animals at different temperatures does not provoke a statistical difference in synaptic sizes. Indeed, at all temperatures the *atg1* mutant animals’ synapses show a synaptic growth comparable to that observed in 15°C control animals (Fig.2 F). We then asked whether Is and Ib boutons were differently affected in the atg1 mutants. We found that the atg1 mutant Is synapses are temperature-independent, showing the same undergrowth phenotype at all temperatures (15°C bouton number is 17.4 ± 2.2; at 25°C it is 19.6 ± 3.7; at 29°C it is 15.9 ± 3.7 (Fig.2 C, D, G). This undergrowth phenotype is also accompanied by Is synapses failing to form. While this is not observed at 15°C, it occurs at 25°C (9% of observed synapses; n = 11) and 29°C (22% of observed synapses; n = 9), illustrating the important function of autophagy at higher temperatures. We also asked whether the temperature-independent Ib boutons were affected by the *atg1* mutation. The control and the *atg1* mutant synapses at 15°C and 25°C presented identical synaptic sizes (Fig. 2H). Surprisingly, Ib synapses in *atg1* mutant animals raised at 29°C were significantly bigger than controls and the *atg1* mutant Ib synapses observed at 15°C and 25°C (Fig. 2 H). We hypothesize that this overgrowth could be a compensatory mechanism of the Ib motoneurons, as a consequence of the severe undergrowth or absence of Is synapses observed at 29°C in *atg1* mutants. This Ib morphological compensation when Is synapses are ablated was also observed in a recent study (Y. Wang et al., 2021). This illustrates another difference between the Is and Ib motoneurons. Indeed, autophagy function can have a drastic effect on the temperature sensitivity of a motoneuron while having a completely different, or no effect on another motoneuron innervating the same muscle. This also raises the interesting possibility that the differences in synaptic growth observed at different temperatures could be a consequence of a difference in autophagic activity at these temperatures.

**Figure 2:**
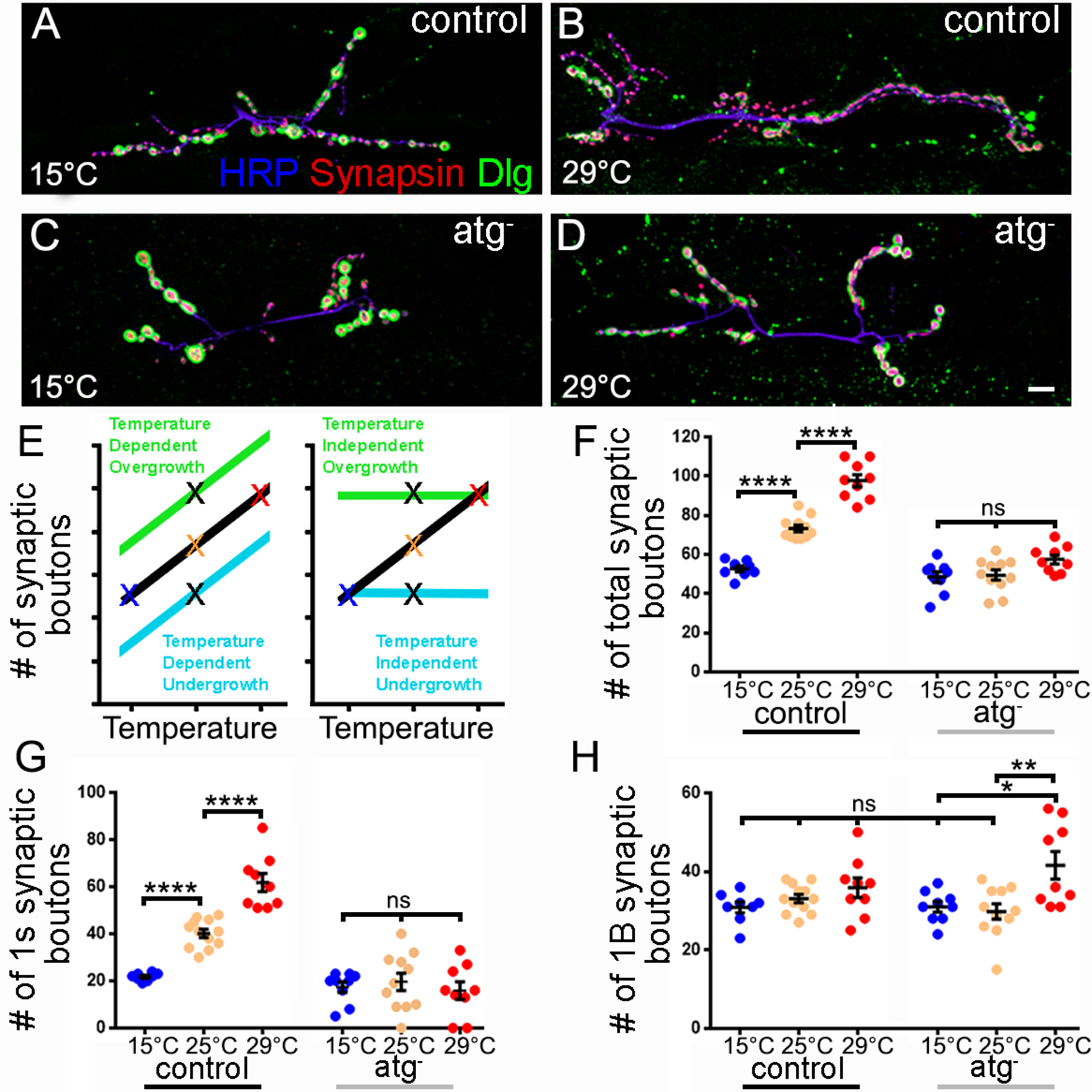
Autophagy is necessary to the temperature dependence of synaptic growth. A-D : Representative NMJs at m6/7 segment A3 in animals reared at 15°C or 29°C with immunofluorescence for a presynaptic membrane marker (blue; anti-HRP), the vesicle marker (red; anti-Synapsin) and the post synaptic marker Discs-Large (green; anti-DLG). E: Schematic diagram depicting the synaptic growth quantification of a temperature-dependent (left) and temperature-independent (right) mutant. The black lines represent the temperature dependent synaptic growth observed. The black “X” represents value from previously characterized overgrown or undergrown mutants. F-H: Quantification of the total, Is and Ib synaptic boutons number at different temperatures in control and atg mutant animals. **** is p < 0.0001, ** is p < 0.01, * is p < 0.05, ns is not significant. ANOVA with post hoc Tukey’s multiple comparisons test. All quantifications show SEM. Scale is 10 μm.

### Autophagy levels change in a temperature-dependent way within the larval CNS

We then investigated whether the level of autophagic activity could vary with rearing temperature. Increases in temperature have been shown to increase the metabolic rate of *Drosophila melanogaster* (Mołoń et al., 2020). Often, an increase in metabolic rate requires an increase in autophagy activity (Rabinowitz & White, 2010). We turned our attention to the protein marker Ref(2)P. This protein, a homolog of P62, is a hallmark marker for autophagy activity, and its function is to flag ubiquitinated substrates for degradation during autophagy (Nezis et al., 2008; Pankiv et al., 2007). As the Ref(2)P protein is degraded along with tagged substrates, accumulation of Ref(2)P indicates impairment of autophagic activity (Bartlett et al., 2011; Nezis et al., 2008). Because this marker had not been tested on *Drosophila* larvae, we decided to observe the immunofluorescence of the Ref(2)P animals in control and *atg1* mutant larval CNS (Fig 3 A and B) as well as at the NMJ (Fig 3 C and D). As expected, the *atg1* mutant animals show stronger immunofluorescence. This is reminiscent of the increase of Ref(2)P protein aggregates observed in the Drosophila adult thoracic muscles, CNS, and fat body cells of autophagy loss of function mutant animals (Nagy et al., 2014; Nezis et al., 2008; Wen et al., 2017). Because Ref(2)P and the Lysotracker staining were ubiquitous it was technically challenging to quantify these markers for autophagy at the NMJ and/or within motoneuron cell bodies. We therefore resorted to a more general quantification of these markers within the brain lobes of the animals, as previously published (Shen & Ganetzky, 2009). In addition, because developmental autophagy is activated in the wandering stage of the 3rd instar larvae (Butterworth et al., 1988; Lee et al., 2002; Rusten et al., 2004), we focused our efforts on the CNS of 2nd instar larvae and asked whether there was a difference in the Ref(2)P intensity between animals raised at 15°C and animals raised at 29°C (Fig 3 E, F and I) . We find that there is a reduction in the Ref(2)P intensity in animals raised at 29°C (intensity is 72.3 ± 9.6 % of the intensity of animals raised at 15°C; p = 0.04). These data suggest that autophagic activity is higher at 29°C than at 15°C. To confirm this possibility, we decided to run a second test to assess autophagy levels. We used the acidophilic dye Lysotracker, which has been used as an indicator of autophagic activity because it labels acidic structures, such as lysosomes (Berry & Baehrecke, 2007; Scott et al., 2007). It has been used to study starvation, cell death, hypoxia, oxidative stress, and for the overexpression of constructs that lead to increased autophagic activity requiring an increase of lysosomes and more acidic structures (Berry & Baehrecke, 2007; Chakraborty et al., 2019; Lai et al., 2016; Scott et al., 2007; Shen & Ganetzky, 2009). We found that Lysotracker levels are significantly increased: at 29°C, intensity is 321.4 ± 69.1 % (n = 11) of the ΔF/F intensity value observed at 15°C (p = 0.001; n = 11). Taken together these results indicate that there is an increase of autophagy correlating with an increase in temperature. We could not distinguish whether autophagy was increased in all or a subset of neurons. Nevertheless, our results suggest that it is the temperature-driven change in autophagy that is at the origin of the Is synaptic increase. We wondered whether we could identify the molecular pathway downstream of the autophagy genes directing the temperature-dependence of synaptic growth.

**Figure 3:**
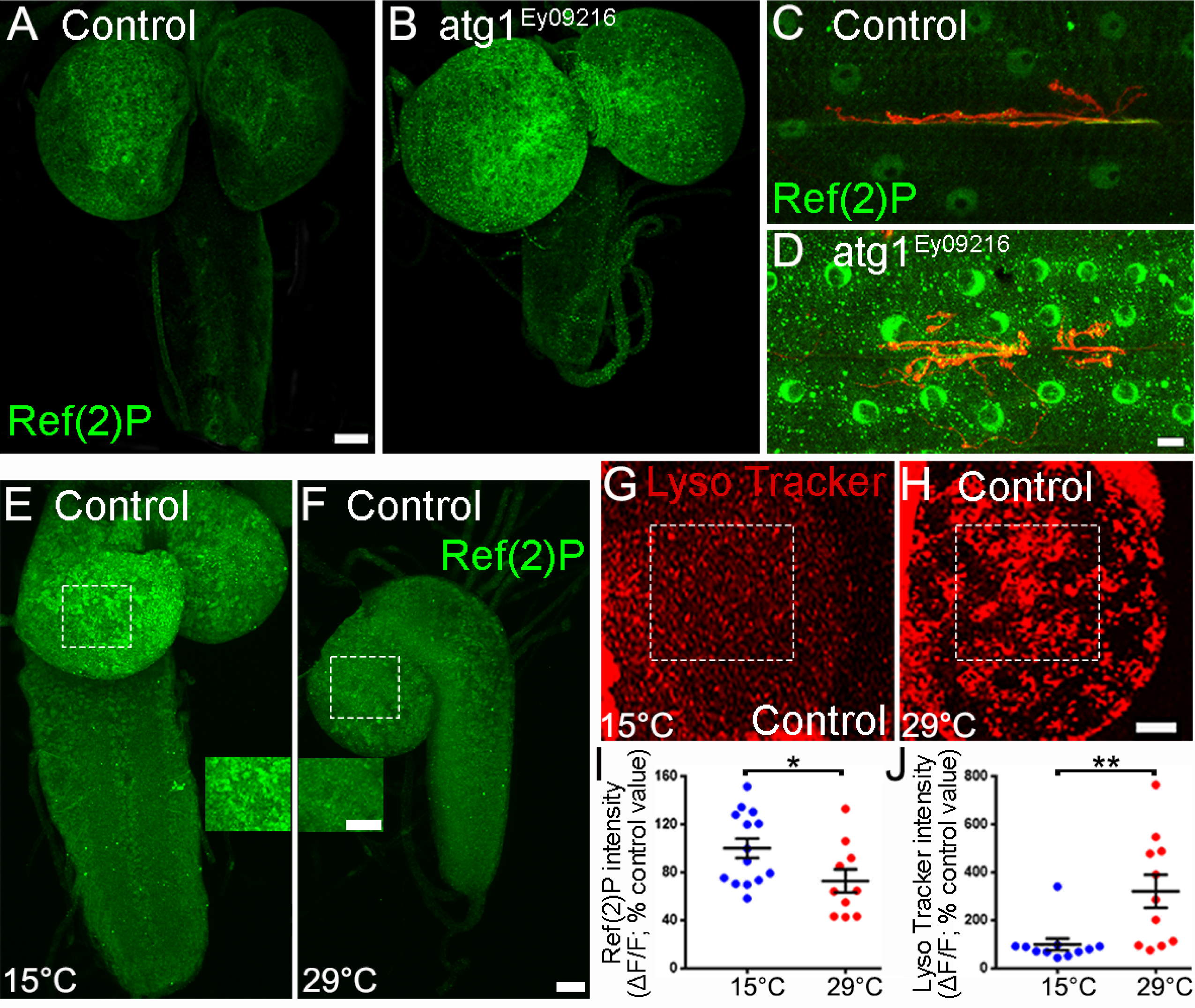
Autophagy activity increases with the rearing temperature. A, B: Representative confocal photograph of a third instar larval CNS showing immunoreactivity for REF(2)P in control and atg mutant animals. C, D: Ref(2)P immunoreactivity is also increased in atg mutant muscles and NMJ (Anti HRP in red). E, F: Representative confocal photograph of a second instar larval CNS showing immunoreactivity for REF(2)P in control animals reared at 15°C and 29°C. Insets show the blow up of a portion of the optic lobes similar to the ones used for quantification purposes. G, H: Representative confocal photograph of a second instar larval optic lobe from control animals raised at 15°C and 29°C showing Lysotracker reactivity. I: Quantification of the Ref(2)P fluorescence intensity in control animals raised at 15°C and 29°C. J: Quantification of the Lysotracker fluorescence intensity in control animals raised at 15°C and 29°C. ** is p < 0.01; * is p < 0.05. An unpaired, two-tailed t test was performed in I. A Mann-Whitney analysis was performed in J. All quantifications show SEM. Scale is 20 μm for A, B, C, D and 10 μm for E, F, G and H.

### The ubiquitin ligase Highwire (Hiw) affects the temperature dependence of synaptic growth

We decided to test the role of the E3 ubiquitin ligase Hiw, the homolog to PAM in humans and RPM-1 in nematodes (Schaefer et al., 2000; Wan et al., 2000). Hiw is a well characterized negative regulator of synaptic growth (Collins et al., 2006; DiAntonio et al., 2001; Wan et al., 2000). Indeed, *hiw* mutants show synaptic overgrowth and Hiw has been shown to be downstream of the autophagy signaling pathway (Shen & Ganetzky, 2009). In fact, *atg* loss-of-function mutants have synaptic undergrowth, which is dependent on Hiw function. The double mutant (*atg, hiw*) shows synaptic overgrowth, resembling that of a *hiw* mutant, while overexpression of *hiw* is sufficient to suppress the synaptic overgrowth phenotype of an *atg* gain-of-function animal. This strongly suggests that *atg* genes exert their function on synaptic growth through repression of Hiw activity (Shen & Ganetzky, 2009).

We asked whether *hiw* mutant synapses could perform temperature-dependent synaptic growth. We noted that the previously described overgrowth occurs similarly at all temperatures. Indeed, *hiw* mutant animals raised at 15°C showed overgrown synapses (102.6 ± 2.3 mean bouton number, n = 18, p < 0.0001) compared to controls raised at the same temperature (53.4 ± 1.4 mean bouton number, n = 19) but this overgrowth was similar to the one observed in *hiw* animals raised at 25°C or 29°C (mean bouton numbers are 104.1 ± 2.4 and 100.4 ± 3.3; p = 0.99 and p = 0.98; Fig 4 A-E). There is no difference in synaptic size between *hiw* mutant animals raised at 15°C, 25°C and 29°C (Fig 4 E). This temperature-independent overgrowth is visible in the Is boutons (Fig 4 F) that are rendered temperature-insensitive in the *hiw* mutant. Indeed, Is boutons show an increase in mean bouton numbers in *hiw* mutants compared to controls (Fig4 F). They have 50.6 ± 2.1 at 15°C (compared to 20.5 ± 1.2 in control; p < 0.0001), 50.2 ± 2.2 at 25°C (compared to 40.2 ± 1.7 in control; p = 0.006) and 52.4 ± 1.7 at 29°C (compared to 61.6 ± 1.5 in control; p= 0.04). In *hiw* mutants there is no statistical difference between synaptic sizes at all temperatures. This is consistent with the hypothesis positing that Hiw is downstream of the autophagic signal and regulates temperature-dependent synaptic growth. In this model animals reared at 15°C present low autophagy, high Hiw synaptic growth-repressive function, and hence small synapses. At 29°C, autophagy is higher, Hiw activity is downregulated, and synapses are larger.

**Figure 4:**
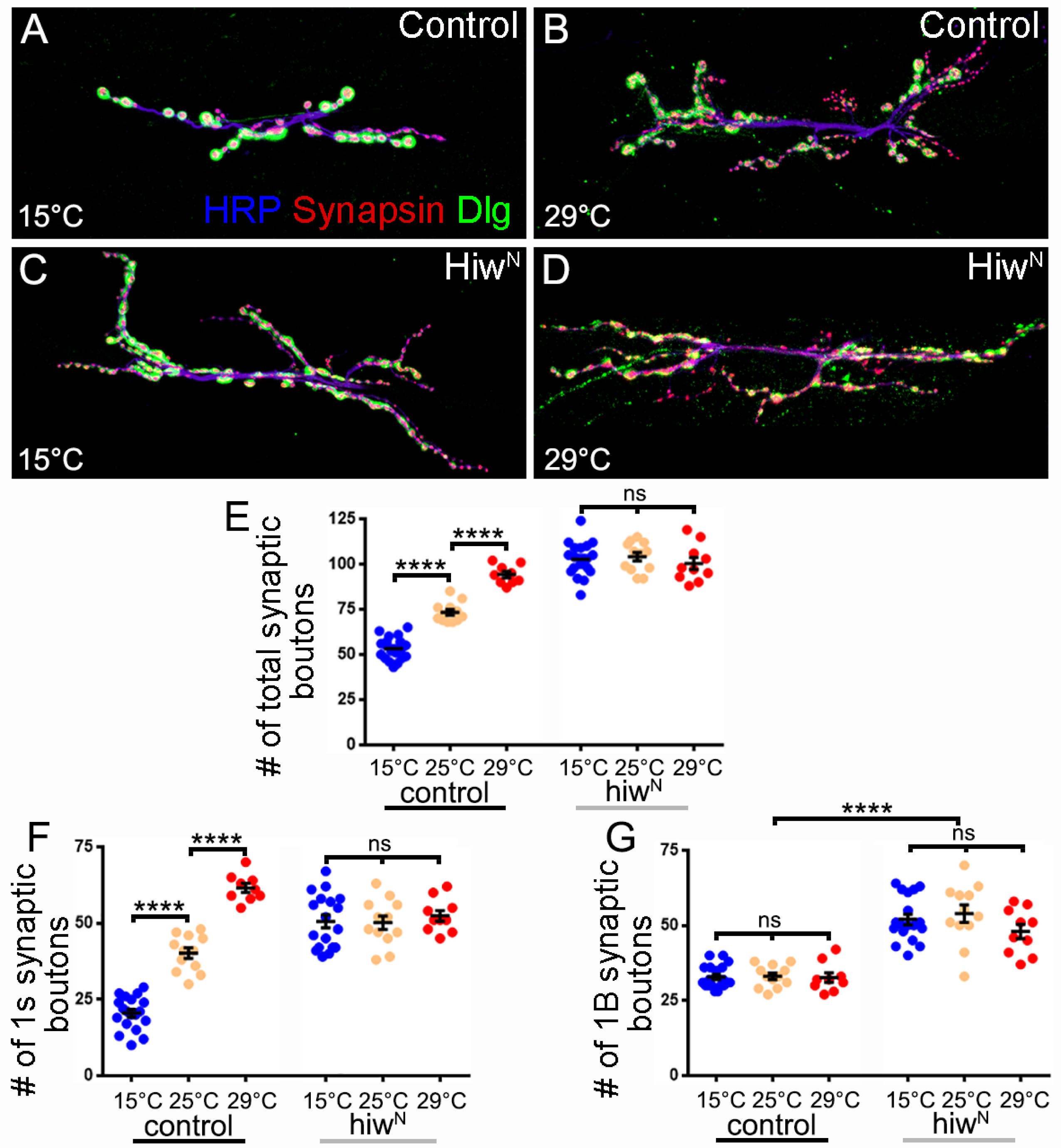
**Hiw affects both temperature -independent and -dependent synaptic growth.**A-D: Representative NMJs at m6/7 segment A3 of control and hiw mutant animals reared at 15°C or 29°C with immunofluorescence for a presynaptic membrane marker (blue; anti-HRP), the vesicle marker (red; anti-Synapsin) and the post synaptic marker Discs-Large (green; anti- DLG). E-G: Quantification of the total, Is and Ib synaptic bouton numbers at different temperatures in control and hiw mutant animals. **** is p < 0.0001 ns is not significant. ANOVA with post hoc Tukey’s multiple comparisons test. All quantifications show SEM. Scale is 10 μm.

Interestingly, *hiw* mutants also show an increase in the mean Ib synaptic bouton numbers (52 ± 1.8 at 15°C; 53.9 ± 2.9 at 25°C and 48 ± 2.4 at 29°C compared to 32.8 ± 0.9, 33.1 ± 1.1 and 32.7 ± 1.6 in control; Fig 4G). Because Ib terminals are temperature-insensitive, we hypothesize that Hiw has a broader function as a negative regulator of synaptic growth in all motoneurons not only the ones influenced by the temperature/autophagy signal.

### The MAPKKK Wallenda (Wnd) controls the increase in synaptic growth taking place at higher temperature (29°C)

We decided to continue investigating the pathway downstream of Hiw. One characterized downstream target of Hiw is the MAPKKK Wallenda (Wnd) (Collins et al., 2006; Wu et al., 2007; Xiong et al., 2010). Wnd and its mammalian ortholog, dual leucine zipper kinase (DLK), are important regulators of neuronal stress pathways (Asghari Adib et al., 2018; Gerdts et al., 2016; Simon & Watkins, 2018). Wnd’s role in axonal regeneration after injury has been extensively studied and it is described as being essential for promoting growth after injury (Hammarlund et al., 2009; Miller et al., 2009; Xiong et al., 2012). Interestingly, Hiw plays a role in regulating Wnd levels in the Drosophila model of Wallerian degeneration. Hiw loss-of-function mutants show an increase in Wnd levels, that in turn leads to a neuroprotective effect in damaged axons (Collins et al., 2006; Lewcock et al., 2007; Nakata et al., 2005; Xiong et al., 2010, 2012). To this day, the role of Wnd in regulating synaptic size remains unclear. Indeed, *wnd* loss-of-function mutations can decrease the synaptic overgrowth observed in *hiw* mutants. This shows that Wnd can promote synaptic growth and is inhibited by Hiw (Collins et al., 2006; Wu et al., 2007; Xiong et al., 2010). Nevertheless, animals only deficient for Wnd showed normal synaptic size, which might indicate that Hiw targets proteins other than Wnd to inhibit synaptic growth. In addition, this could mean that a different stimulus is required to recruit Wnd to promote synaptic growth (Collins et al., 2006). It is important to keep in mind that these experiments were typically carried out at room temperature (about 22°C) or at 25°C. Here, we decided to investigate the role of Wnd on temperature-dependent synaptic growth. As previously, we carried out immunohistochemistry to analyze the number of synaptic boutons in control and wnd mutant animals raised at 15°C, 25°C and 29°C. We observed unaffected temperature-dependent synaptic growth from 15°C to 25°C in both control and *wnd* mutants (Fig 5 E). At 15°C, synapses show a mean bouton number of 52.6 ± 1.5 (control; n = 16) and 54 ± 2.4 (*wnd*; n = 9; p = 0.99) while at 25°C, we observed 73.3 ± 1.6 (control; n = 12) and 73.7 ± 3 (*wnd*; n = 12; p = 0.99). This is consistent with the previous studies showing that *wnd* mutations did not impact synaptic growth (Collins et al., 2006; Shen & Ganetzky, 2009). The difference between control and *wnd* mutant synapses was obvious at higher temperature. While control animals showed an increase in synaptic size at 29°C compared to 25°C (92 ± 4.4; n = 8; p < 0.0001), *wnd* mutant animals at 29°C presented a synapse with a similar size (73.7 ± 2; n = 13) to the one observed at 25°C (p > 0.99). This illustrates a role for Wnd in promoting synaptic growth at higher temperatures. When assessed in more detail, we noted that control and *wnd* mutant Ib synapses were not sensitive to temperature (Fig 5 G). The absence of supplementary synaptic growth observed at high temperatures in the *wnd* mutants was totally due to the absence of supplementary synaptic growth of the Is synapse. At 15°C and 25°C the growth of the Is synapse was similar in control (20.9 ± 1 and 40.2 ± 1.7) and *wnd* mutant animals (23.2 ± 2.8 and 39.1 ± 2; Fig 5F), showing a normal temperature sensitivity. The data collected at 29°C was very different. While controls still showed temperature-sensitive synaptic growth (at 29°C synaptic growth increased to 56.9 ± 3.2 mean boutons number; p < 0.0001), *wnd* mutant animals show no growth between 25°C and 29°C (compare 39.1 ± 2 at 25°C to 37.5 ± 2 at 29°C; p = 0.99). This shows that the MAPKKK Wnd is specifically involved in regulating synaptic growth in a subset of motoneurons.

**Figure 5:**
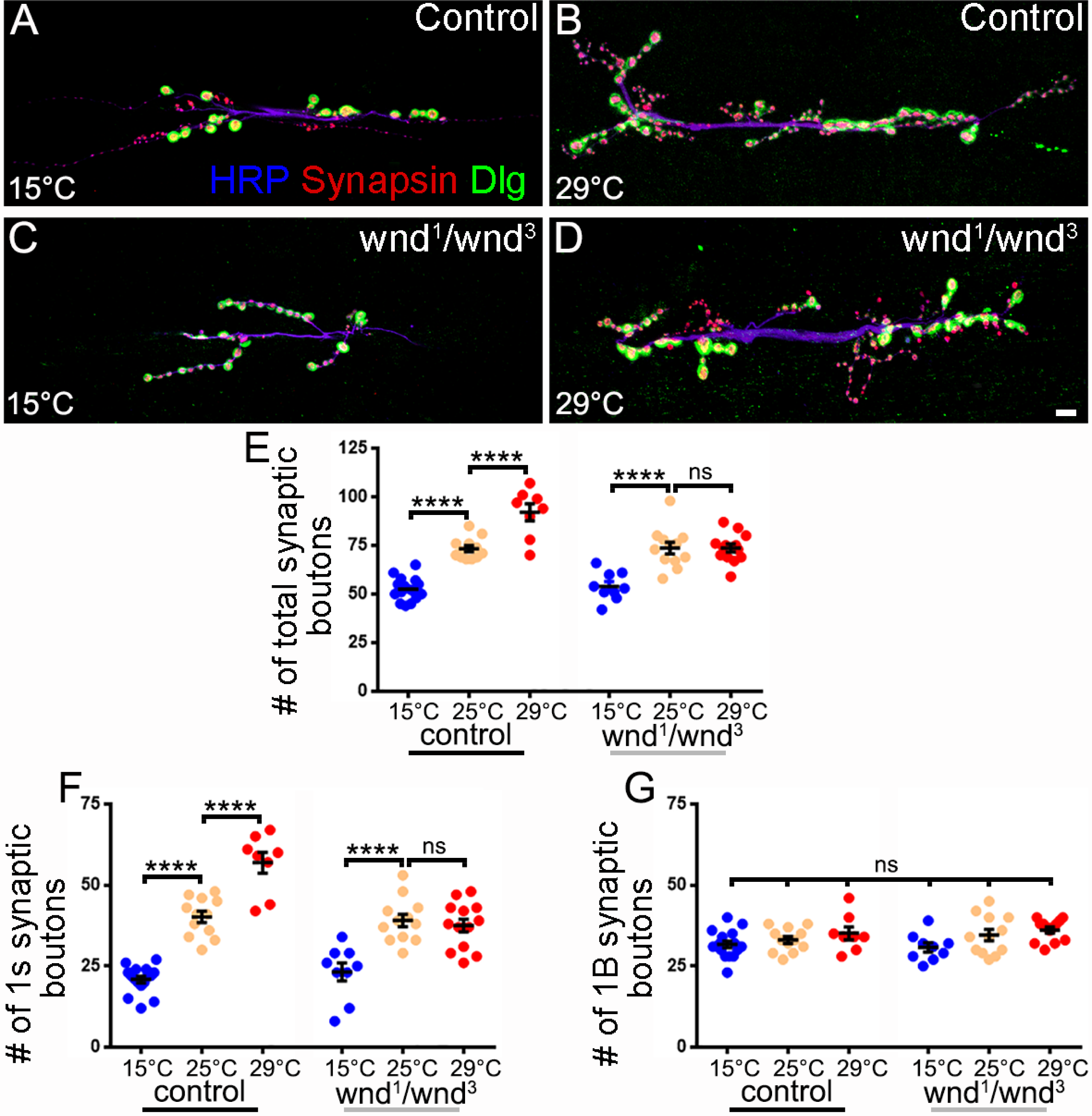
The MAPKKK Wnd is necessary for the temperature dependence of synaptic growth occurring at 29°C. A-D: Representative NMJs at m6/7 segment A3 of control and wnd mutant animals reared at 15°C or 29°C with immunofluorescence for a presynaptic membrane marker (blue; anti-HRP), the vesicle marker (red; anti-Synapsin) and the post synaptic marker Discs-Large (green; anti- DLG). E-G: Quantification of the total, Is, and Ib synaptic bouton numbers at different temperatures in control and wnd mutant animals. **** is p < 0.0001 ns is not significant. ANOVA with post hoc Tukey’s multiple comparisons test. All quantifications show SEM. Scale is 10 μm.

### The MAPK P38b functions downstream of Wnd to control synaptic growth at higher temperature (29°C)

Having discovered a new function for the MAPKKK Wnd, we decided to investigate the downstream molecular players in this pathway. There are 2 possible MAPKK that can be activated by Wnd, the Mkk7 and Mkk4 homologs known as Hemipterous (Hep) and Mkk4 (Han et al., 1998; Holland et al., 1997; Thibault et al., 2004). We used the loss of function alleles *hep^r75^* and *mkk4^e01485^* because *Hep^r75^*has been documented to be lethal at the pupal stage (Bosch et al., 2005), while *mkk4* loss-of-function animals appeared viable under normal rearing conditions (Hsu et al., 2021). Nevertheless, when we reared these mutants at 29°C, we found that lethality was occurring earlier than previously described, and we were unable to isolate third instar larvae to assess our model NMJ. This suggests that both MAPKK may have a distinctive role in animals raised at 29°C. We decided to ask whether we could isolate a downstream target of these MAPKK. We found that the inhibition of Jun Kinase by the overexpression of RNAi constructs in neurons and that *p38a* null mutant animals did not affect the NMJ growth of animals reared at 29°C (data not shown). We found that a loss-of-function mutation in the MAPK *p38b* phenocopies the *wnd* mutant phenotype. Indeed, *p38b* mutants do not show any differences from control when we consider the mean number of synaptic boutons at 25°C (Fig 6 A, C, E; mean bouton number is 74.8 ± 2 in control and 71.9 ± 1.3 in *p38b* mutants; p = 0.7). We observe a mean Is bouton number of 41.7 ± 1.8 in control and 39.7 ± 1.2 in *p38b* mutants (Fig 6 E; p = 0.81) and mean Ib boutons number of 33.1 ± 0.8 in control and 32.2 ± 1 in *p38b* mutants (Fig 6 F; p = 0.96). In contrast, there is a clear difference when we compare the mean bouton number at 29°C. While control preparations show temperature- dependent growth (total bouton number is 94.2 ± 1.7 at 29°C, an increase compared to the 25°C value; p <0.0001), *p38b* mutants show no additional growth at 29°C (mean boutons number 69.2 ± 2; p = 0.68) when compared to the synaptic growth at 25°C (71.9 ± 1.3). As before there is no difference in the Ib synapses at 29°C (Fig 6 G) but surprisingly, like in *wnd* mutants, *p38b* mutants did not show a Is synaptic bouton increase at 29°C (39.7 ± 1.2 at 25°C and 35.7 ± 2 at 29°C; p = 0.25; fig 6 F). This strongly suggests that P38b is the MAPK downstream of the MAPKKK Wnd responsible for the additional synaptic growth observed at 29°C. To further test this hypothesis, we carried out a genetic interaction test (Fig 6 H). We tested the synapses for their ability to grow at 29°C. In this condition, control animals (mean bouton number 96.5 ± 2), *wnd/+* heterozygote animals (mean bouton number 91.6 ± 2.6) and *p38b/+* heterozygote animals (mean bouton number 93.6 ± 1.8) grew to a very similar size. This is not surprising since most genes do not show haploinsufficiency [for example only 300 human genes show haploinsufficiency; (Dang et al., 2008; Deutschbauer et al., 2005)]. In contrast, when the animal contains 50% of Wnd and 50% of P38b (*p38b/+; wnd/+* double heterozygotes) we observed a lower mean bouton number (76.2 ± 2.6). This is a statistically significant decrease compared to controls, or to animals carrying a single heterozygote mutation (Fig 6 H), and is a strong indication that Wnd and P38b belong to the same pathway.

**Figure 6:**
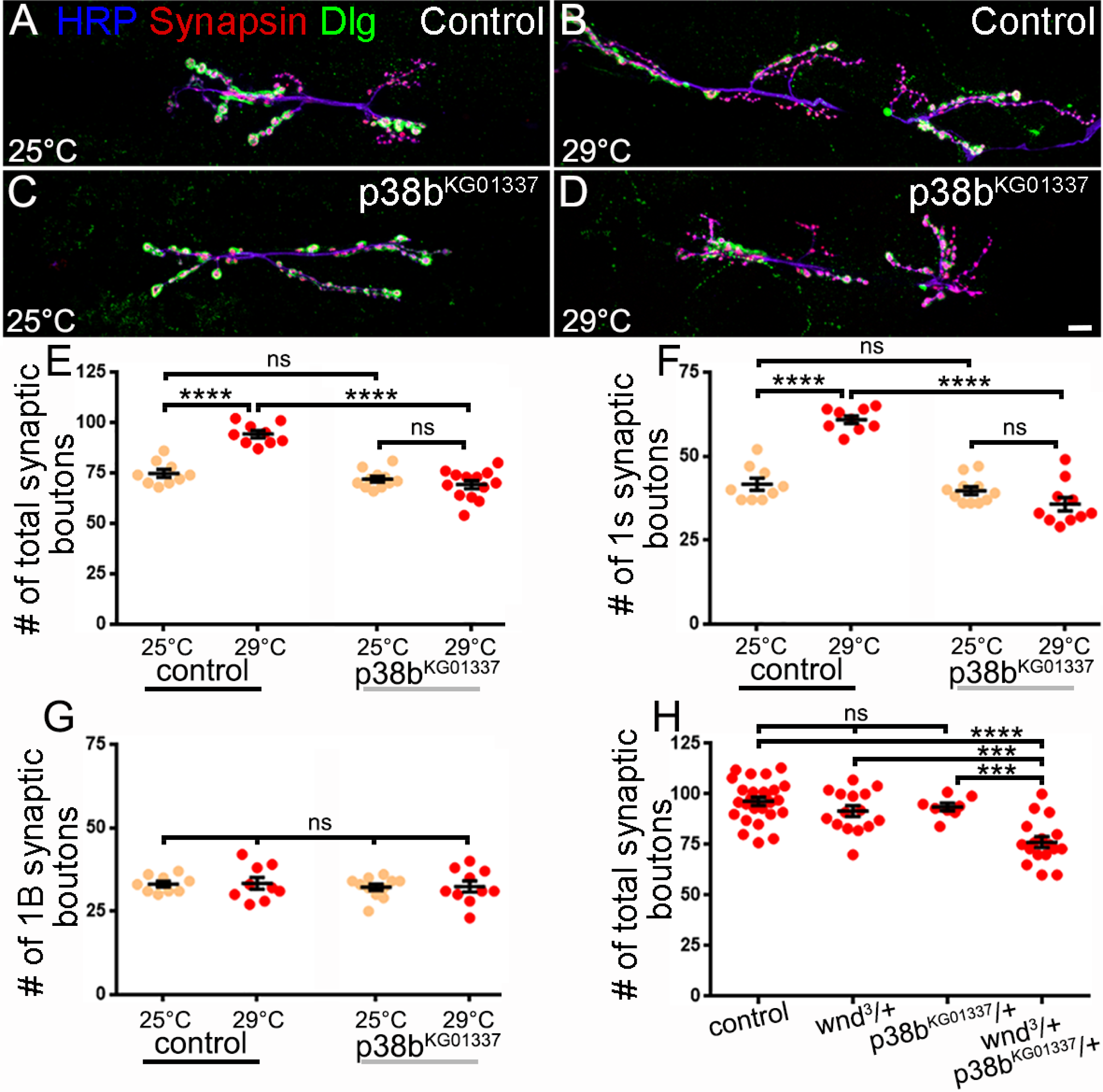
The MAPK P38b is necessary to the temperature dependence of synaptic growth occurring at 29°C and interacts with wnd. A-D: Representative NMJs at m6/7 segment A3 of control and p38b mutant animals reared at 25°C or 29°C with immunofluorescence for a presynaptic membrane marker (blue; anti-HRP), the vesicle marker (red; anti-Synapsin) and the post synaptic marker Discs-Large (green; anti- DLG). E-G: Quantification of the total, Is, and Ib synaptic boutons number at 25°C and 29°C in control and *p38b* mutant animals. H: Quantification of the total synaptic boutons number at 29°C of control, heterozygote animal *wnd/+* and *p38b/+* and the transheterozygote *wnd/+; p38b/+*. **** is p < 0.0001, *** is p < 0.001, ns is not significant. ANOVA with post hoc Tukey’s multiple comparisons test. All quantifications show SEM. Scale is 10 μm.

Taken together these data show that there is a temperature-dependence of synaptic growth specific to a subset of motoneurons and regulated by a molecular pathway including the *atg*, *hiw*, *wnd* and *p38b* genes.

## Discussion

Temperature can impact cellular properties, and cold- or heat-shock responses have been well documented (Al-Fageeh & Smales, 2006; Knapp & Huang, 2022; Parsell & Lindquist, 1993; Phadtare et al., 1999; Velazquez & Lindquist, 1984). Nevertheless, more subtle variations in temperature that do not provoke protein damage can also have an impact on neuronal properties. In vertebrate, axonal conduction, action potential shape, and neurotransmitter release can vary with naturally occurring subtle changes in temperature (2°C to 3°C; Andersen & Moser, 1995). Optogenetic experiments in mice showed that fluctuations of 0.2 to 2°C can suppress spiking activity in multiple brain regions (Owen et al., 2019). In *Drosophila*, animals raised at 28°C instead of 18°C showed an increase in one specific voltage-gated potassium channel, consequently affecting the action potential duration and amplitude (Chopra & Singh, 1994). Here, we characterize the effect of varying temperature across a span of 14°C ranging from 15°C to 29°C and show that this difference in temperature can have spectacular consequences on synaptic growth. Indeed, the temperature-driven increase in synaptic material is not subtle; the Is synapse shows a 200% increase in size. Because the Is neuron is a phasic bursting neuron innervating multiple muscles on each hemi-segment, it is tempting to hypothesize that the increase of its synaptic size with temperature is linked with the observed increased of motility at higher temperatures (Lnenicka & Keshishian, 2000; Schuster et al., 1996; Sigrist et al., 2003). Interestingly, when there is a reduction in the Is input, muscle contraction remains intact, while there is an increase in the failure of waves that propagate through multiple segments (Y. Wang et al., 2021). These observations reinforce the view that tonic Ib motoneurons act as primary drivers of contraction, while phasic Is motoneurons coordinate the contraction of muscle subgroups and direction (Heckscher et al., 2012; Hoang & Chiba, 2001; Johansen et al., 1989; Newman et al., 2017; J. E. Schaefer et al., 2010).

Another remarkable feature of the temperature-dependent synaptic growth is that it is specific to a subset of motoneurons. Indeed, the synaptic growth of the Ib motoneuron that innervates the same target muscle as the Is NMJ is unaffected by temperature. This last point reinforces the growing literature demonstrating the differences between the tonic Ib and the phasic Is motoneurons. In short, Ib motoneurons have a lower probability of release, undergo facilitation and sustained release during stimulation, while Is motoneurons show higher probability of release and rapid depression during high frequency stimulation (Aponte-Santiago et al., 2020; Lu et al., 2016; Newman et al., 2017; Nguyen & Stewart, 2016; Y. Wang et al., 2021). Interestingly, a recent study has recently characterized genes that are differentially expressed in Is and IB motoneurons, which may be the molecular basis of these differences (Jetti et al., 2023); in particular they mention that Atg1 and Atg4a mRNAs are enriched in Is neurons. We have now determined that the temperature dependence of synaptic growth and its molecular regulation by autophagy genes and the MAPK pathway is only relevant to 1s synapses. It is not due to the inability of the Ib synapse to be plastic. Indeed, although insensitive to temperature, the Ib motoneuron is capable of structural plasticity. When the Is motoneuron activity is reduced, the Ib motoneuron can compensate by increasing its synaptic size and neurotransmitter release (Aponte-Santiago et al., 2020; Y. Wang et al., 2021). The Is motoneuron remains largely insensitive to changes in Ib input manipulation (Aponte-Santiago et al., 2020; Y. Wang et al., 2021).

Our results strongly point to autophagy levels being a master regulator of temperature-dependent synaptic growth. We have used two different methods (Ref2(P)/P62 level of expression and the Lysotracker levels of reactivity) to show that autophagy was increased within the CNS of *Drosophila* raised at higher temperature. It is important to stress that the previously reported synaptic undergrowth associated with the autophagy deficiency (Shen & Ganetzky, 2009) is solely due to the contribution of the phasic Is synapses and that the Ib synapse is greatly unaffected at 15°C and 25°C, only showing a marginal increase at 29°C; an increase that could be a compensatory phenomenon to the drastic reduction or absence of the Is synapses. Interestingly, the autophagy signal was previously shown to be a positive regulator of synaptic growth by inhibiting the ubiquitin ligase and negative regulator of synaptic growth Hiw (Shen & Ganetzky, 2009). It is tempting to draw a model in which elevated temperature provokes elevated autophagy activity which in turn causes lower Hiw activity leading to bigger synapses. Our results also confirm that Hiw is a master controller of synaptic growth (Collins et al., 2006; Wan et al., 2000), involved in regulating both temperature-dependent and - independent synaptic growth. The MAPK pathway is one of the signaling pathways regulated by Hiw and involved in synaptic growth. A very potent argument is that the overgrown synapses encountered in the *hiw* mutant are restored towards control values in the *hiw, wnd* double mutant (Collins et al., 2006). Because the over expression of Wnd can provoke overgrowth (Collins et al., 2006), the working hypothesis was that Wnd signaling cascade was the driving force beyond synaptic growth, controlled and repressed by Hiw (Collins et al., 2006; Klinedinst et al., 2013). Nevertheless, experiments carried out at room temperature or 25°C did not show a decrease in synaptic growth in the *wnd* mutant.

Here our experiments reveal a function for Wnd at 29°C supporting the fact that it is responsible for the additional growth seen at higher temperature. We propose that the higher autophagy activity at 29°C can sufficiently repress Hiw function to reveal the function of Wnd in synaptic growth. This last point may illustrate the fact that research is carried out in strict and narrow laboratory conditions and changing one environmental parameter (like temperature) might be used as a tool to reveal new functions.

## Acknowledgments

We thank Dr. Jonathan Blagburn for his valuable comments on previous versions of this manuscript.

